# A response-locked classification image analysis of the perceptual decision making : contrast detection

**DOI:** 10.1101/2021.06.17.447463

**Authors:** Hironori Maruyama, Natsuki Ueno, Isamu Motoyoshi

## Abstract

In many situations, humans make decisions based on serially sampled information through the observation of visual stimuli. To quantify the critical information used by the observer in such dynamic decision making, we here applied a classification image (CI) analysis locked to the observer’s reaction time (RT) in a simple detection task for a luminance target that gradually appeared in dynamic noise. We found that the response-locked CI shows a spatiotemporally biphasic weighting profile that peaked about 300 ms before the response, but this profile substantially varied depending on RT; positive weights dominated at short RTs and negative weights at long RTs. We show that these diverse results are explained by a simple perceptual decision mechanism that accumulates the output of the perceptual process as modelled by a spatiotemporal contrast detector. We discuss possible applications and the limitations of the response-locked CI analysis.

## 1. Introduction

While humans and animals can immediately recognize objects and scenes at a glance (Thorpe, Fize & Marlot, 1996; Motoyoshi, Nishida, Sharan & Adelson, 2007; Whitney & Yamanashi, 2018), in many situations they ensemble information in a sequence to take more appropriate decisions (Bergen & Julesz 1983; Treisman & Gelade, 1980; Wolfe, 2015). In cognitive psychology, such dynamic information processing has been investigated mainly by measuring the reaction times and correct rates of observers. However, the reaction time alone is not powerful enough to reveal what kind of information in the stimuli led the observers to make a decision at that moment in time, unless data obtained under various conditions are compared.

In visual neuroscience, reverse correlation analysis is widely applied to reveal the information in stimuli that determines the system responses (Neri & Levi, 2006; Neri, Parker & Blakemore, 1999). This analysis has been applied not only to the responses of cortical neurons (DeAngelis, Ohzawa & Freeman, 1993), but also to the analysis of the behavioral responses of human observers (Ringach, 1998; Solomon, 2002). The classification image (CI) method, one such technique, visualizes what information in the stimuli observers consider important for a given perceptual judgement (Ahumada, 1996; Abbey & Eckstein, 2002; Abbey & Eckstein, 2007). In typical experiments, the observer’s responses to a visual target embedded in white noise are collected, and the information in the stimulus that affected the observer’s response is mapped out by analyzing the correlation between the noise and the response in each trial. The CI technique has been used to reveal the spatial distribution of information, or perceptive field, that determines the observer’s judgments for a variety of visual tasks (Eckstein, Shimozaki & Abbey, 2002; Gold, Murray, Bennett & Sekuler, 2000; Rajashekar, Bovik & Cormack, 2006).

With dynamic stimuli, the CI method can also yield spatiotemporal perceptive fields (Neri & Heeger, 2002; Mareschal, Dakin & Bex, 2006; Neri & Levi, 2008). Neri & Heeger (2002) analyzed the correlation between spatiotemporal noise and responses in each trial in a contrast detection task for luminance bars that slowly appear in dynamic noise. They found CI profiles with biphasic weights in time and space, similar to the spatiotemporal impulse response of the early visual system. Recently, similar psychophysical reverse correlation with dynamic stimuli has been applied to the judgement on the average of time-varying visual information to investigate the mechanisms of perceptual decision making (Gardelle & Summerfield, 2011; Hanks & Summerfield, 2017; Li, Castañón, Solomon, Vandormael & Summerfield, 2017; Vandormael, Castañón, Balaguer, Li & Summerfield, 2017; Sato & Motoyoshi, 2020; Summerfield & Tsetsos, 2015; Yashiro & Motoyoshi, 2020; Yashiro, Sato, Oide & Motoyoshi, 2020).

In the aforementioned studies, however, observers made decisions after the visual stimuli had been shown. Such a judgment, which is usually based on visual working memory, is somehow dissociated from on-the-fly judgments that we make in the real world. To clarify when observers make decisions and what information observers rely on to make decisions during observation, one can analyze correlations at each time point of the stimulus locked to the reaction time of the observer during the presentation of the stimulus rather than the stimulus onset. This response-locked reverse correlation has been employed in several studies (Ringach, 1998; Busse, Katzner, Tillmann & Treue, 2008; Caspi, Beutter & Eckstein, 2004; Okazawa, Purcell & Kiani, 2018). For example, Caspi et al. (2004) examined visual features that trigger saccadic eye movements by analyzing the noise at time points locked to the onset of the saccade while the observers views a multi-element display. Okazawa et al. (2018) adopted a reverse correlation analysis locked to button-press responses to stochastic motion to explore the properties of global-motion detectors and decision-making mechanisms.

In the present study, we applied the response-locked CI analysis to the most basic visual task, luminance contrast detection. Specifically, we used stimuli similar to those used by Neri & Heeger (2002) to measure responses and reaction times for target stimuli that emerge slowly in dynamic noise, and we then analyzed the correlation between the noise and response at each time point backward, locked to the observer’s reaction time. This protocol allowed us to examine what signals and what point in the stimulus determined the observer’s decision about the target and the observer’s reaction time. The results revealed spatiotemporally biphasic CIs similar to those reported by Neri & Heeger (2002). On the other hand, we also found that the profile of the CI substantially varied depending on the response time of the observer in a way that was unpredictable from the response properties of the early visual system. These apparently complicated results, however, were quantitatively described by a simple computational model incorporating a perceptual process approximated by a spatiotemporal filter and a decision process (drift-diffusion) that accumulates its output.

## 2. Methods

### 2.1. Observers

Five naïves and two of the authors (average age: 22.8 years) with corrected-to-normal vision participated in the experiment. All experiments were conducted with permission from the Ethics Committee of the University of Tokyo. Observers gave written informed consent. The study followed the Declaration of Helsinki guidelines.

### 2.2. Apparatus

Visual stimuli were displayed on a gamma-corrected LCD monitor (BENQ XL2735) controlled by a PC. The refresh rate was 60 Hz, and the pixel resolution was 0.04 deg/pixel at the viewing distance of 50 cm that we used. The mean luminance of the uniform background was 88.9 cd/m2. All experiments were conducted in a dark room.

### 2.3. Stimuli

The visual stimulus was square dynamic one-dimensional noise (4.8 × 4.8 deg) comprising 16 vertical bars with a width of 0.3 deg (Fig. 1). The contrast (*C_noise_ (t)*) of each bar was switched at a frame rate of 30 Hz according to Gaussian noise with an RMS contrast of 0.1. The total duration was 8000 ms. Two independent 1D-noise fields were presented adjacent to the fixation point.

**Figure 1.**
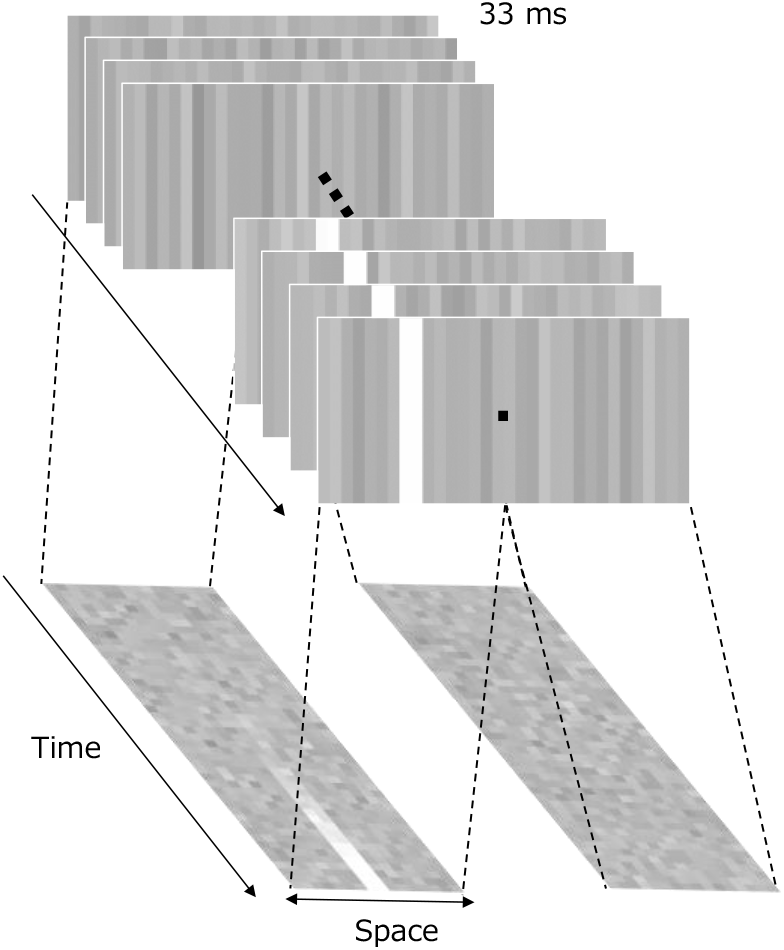
Schematic diagram of the visual stimuli used in the experiment. The bright bar that appears on the left is the target.

The target signal (*C_target_ (t)*) was linearly added to the two central bars only for one of the noise fields. The contrast of the target signal, *C_target_ (t)*, increased linearly with time (*t*) on a logarithmic scale according to the following equation.

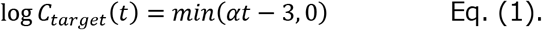

Here, *t* is the time from the stimulus onset. α is the rate of increase, which was set at three levels: 0.05, 0.1, and 0.2. The contrast of each bar was clipped in the range of −1 to +1. The two fields, with and without the target signal, were transformed into luminance images using the relation L(t) = L_mean_ (1+C(t)), where L_mean_ is the mean luminance of the uniform background (88.9 cd/m^2^).

### 2.4. Procedure

In each trial, observers viewed the stimulus at a fixation point binocularly and indicated by pressing a button whether the target appeared in the left or right noise field as quickly as possible. If an observer’s response exceeded the deadline (8000 ms) or was an error, auditory feedback was given, and the data recorded in that trial were excluded from the analyses. The next trial started no less than 0.5 s after the observer’s response. The average error rates were 0.03, 0.02, and 0.02 for contrast increases (α) of 0.05, 0.1, and 0.2, respectively. In each trial, the contrast values of all individual bars (C_noise_ (x,t)), the observer’s response (left, right), and the reaction time were recorded. Each session of the experiment comprised 150 trials for a single condition. For each observer, sessions were repeated until at least 1200 trials were conducted for each condition.

## 3. Results

### 3.1. Reaction time

Figure 2a shows the average logarithmic reaction time of the observer, plotted as a function of the contrast increase (α). On a linear scale, the reaction times were 1949 ms (s.e.= 36.1), 1207 ms (s.e.= 20.7), and 818 ms (s.e.= 22.7) for a contrast increase (α) of 0.05, 0.01 and 0.2, respectively. Fig. 2b is a cumulative histogram of the reaction times of all observers. Fig. 2b shows that a slower rate of increase in the contrast resulted in a longer average reaction time. One-way repeated-measure ANOVA on the average reaction time with increasing contrast showed a significant effect of the rate of contrast increase on the reaction time (F(2, 12) = 1574.6, p < 0.001).

**Figure 2.**
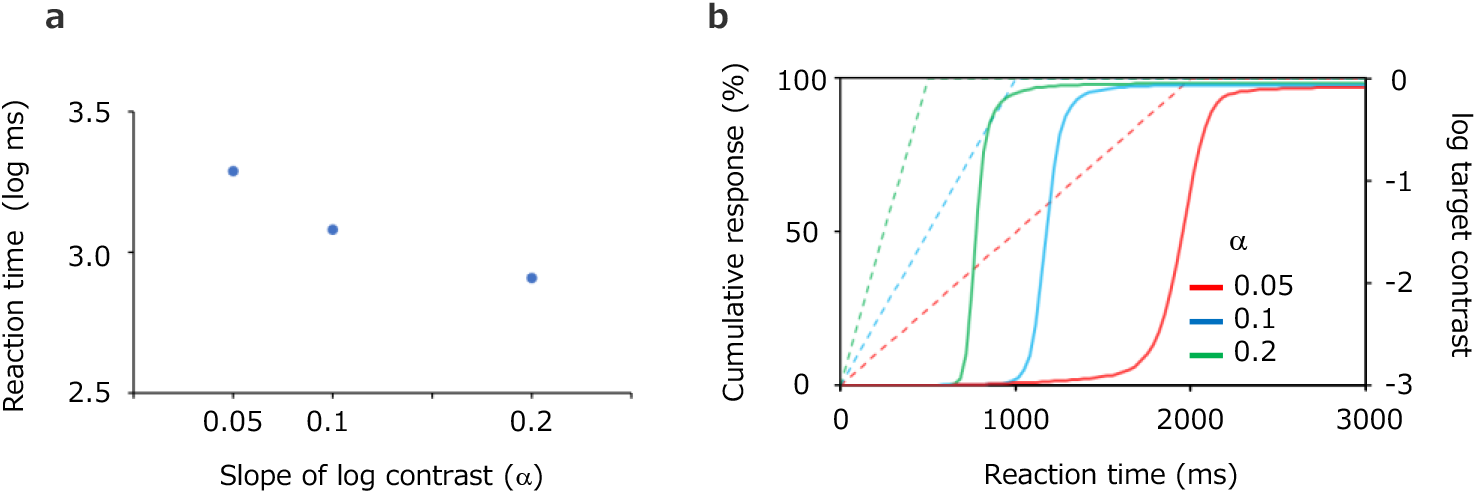
(a) Average logarithmic reaction time as a function of the contrast increase rate. Error bars represent ±s.e. across observers (invisibly small). (b) Cumulative histogram of reaction times of all observers (solid line). Dashed lines represents the logarithmic contrast of the target stimulus over time. Red, blue, and green lines represent a contrast increase of 0.05, 0.1, and 0.2, respectively.

### 3.2. Reverse correlation analysis locked to the response time

We conducted a reverse correlation analysis between the contrast of each bar and the observer’s response (left, right) at each time (*t*) back from the reaction time to characterize the noise common to the time before the reaction. Fig. 3 is a diagram of the analysis. As in Neri & Heeger (2002), μ_1_(x,t) is the mean of the noise contrast in the region where the observer responded that the target was present and μ_0_(x,t) is the mean of the noise contrast in the region where the observer responded that the target was not present. The results were calculated as follows.

**Figure 3.**
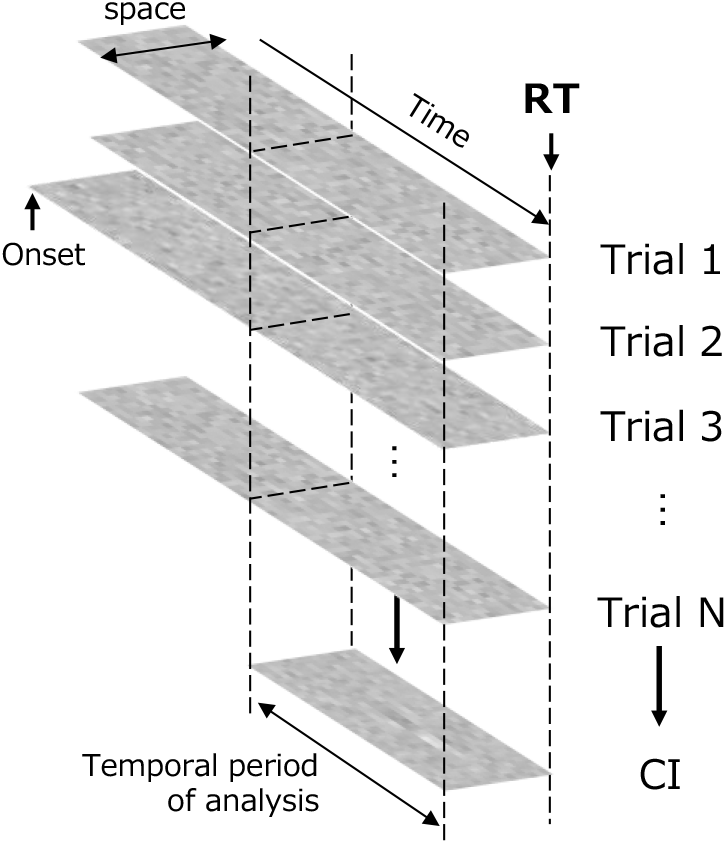
Reverse correlation analysis locked to the reaction. The classification image (CI) was calculated for each bar contrast at each time from the reaction time.

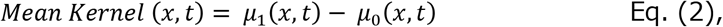

here Mean Kernel refers to the effect of the noise contrast on the response.

The upper panels in Fig. 4 show the classification image (i.e., Mean Kernel) obtained in the reverse correlation analysis locked to the reaction time. The horizontal axis represents the time (*t*) back from the reaction time, and the vertical axis represents the spatial position of each bar (*x*). In comparison with the grey background, the brighter points represent positive weights and the darker points represent negative weights. Individual panels show results for a contrast increase (α) of 0.05, 0.1, or 0.2. The lower panels show the mean of the weights of the two central bars (red) and the mean of the weights of the two surrounding bars adjacent to them (blue) in the CI. The vertical axis represents the weight and the horizontal axis represents the time from the reaction time. We refer to the plots as impact curves.

**Figure 4.**
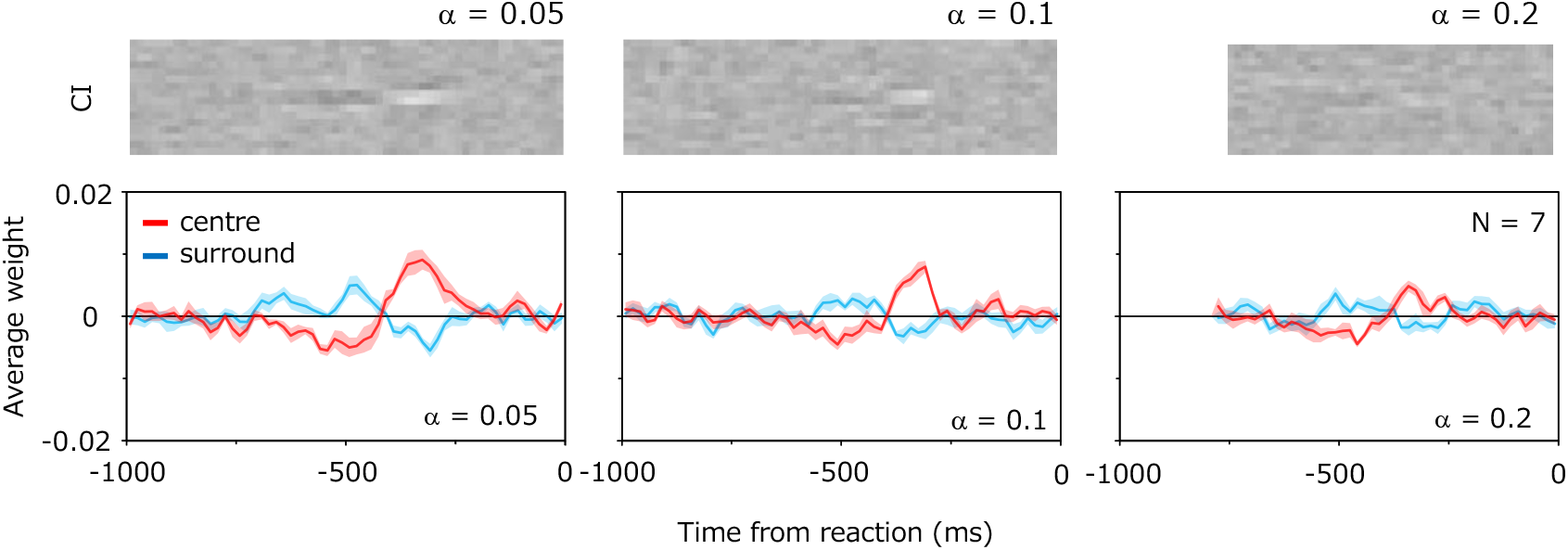
Results of response-locked reverse correlation analysis. The upper panels show the classification image (CI). The vertical axis represents the space and the horizontal axis represents the time from the reaction time. Each pixel represents a positive (bright) or negative (dark) weight. The lower panels show the average of the weights in the central two bars (red curves) and the average of the weights in the two adjacent bars (blue curves). The vertical axis represents the weights and the horizontal axis represents the time from the reaction time. Error bars represent ±s.e. across observers. Individual panels show the results for a contrast increase (α) of 0.05, 0.1, or 0.2.

The above plots show characteristic temporal and spatial variations in the weights before the target detection response. At the center of the stimulus where the target appeared, a large positive weight was found about 300 ms before the response, and a negative weight was found about 500 ms before the response. For the spatial variation, we find that the weights around the target are reversed from the center. This indicates that the central bar in which the target appears has high luminance compared with the surrounding bars, which is a cue to the response. It is also found that the absolute magnitude of the weights tends to increase as the contrast increase rate increases. We also conducted the same analysis for contrast variance, as was done in a previous study (Neri & Heeger, 2002), but found no clear CI profile at all.

For each condition, we calculated the average weights during the time epoch from −250 to −350 ms when in a clear variation in the weights is observed. We then conducted a two-way repeated-measure ANOVA on the average weights with factors of the position (center and periphery) and the contrast increase rate (a = 0.05, 0.1, 0.2) for each condition. The results reveal that the main effect of position (F(1,6) = 182.15, p < 0.001) was significant while the main effect of the contrast increase rate (F(2,12) = 1.689, p = 0.26) was not significant, and there was a significant interaction between the factors (F(2,12) = 20.28, p < 0.001).

### 3.3. Relationship between the RT and CI

As shown in Fig. 2, the reaction time of the observer varied even under the same conditions. Each individual observer responded quickly in some trials and took a long time in others. Taking advantage of this fact, we investigated if and how the CI changes with the reaction time. To this end, we divided the observer’s data into 50% trials with short reaction times, 50% trials with intermediate reaction times, and 50% trials with long reaction times for each condition of the contrast increase rate, and carried out the reverse correlation analysis for each group.

Figure 5 shows the CIs and impact curves obtained for each reaction time group. The results were surprisingly different across the groups. The positive weights about 300 ms before the reaction are larger for the shorter reaction time group. Conversely, the negative weights 500 ms before the reaction are larger for the longer reaction time group. This tendency is constant regardless of the contrast increase rate (α). The results indicate that the spatiotemporal profile of the weights of information correlated with the response is remarkably different depending on the reaction time.

**Figure 5.**
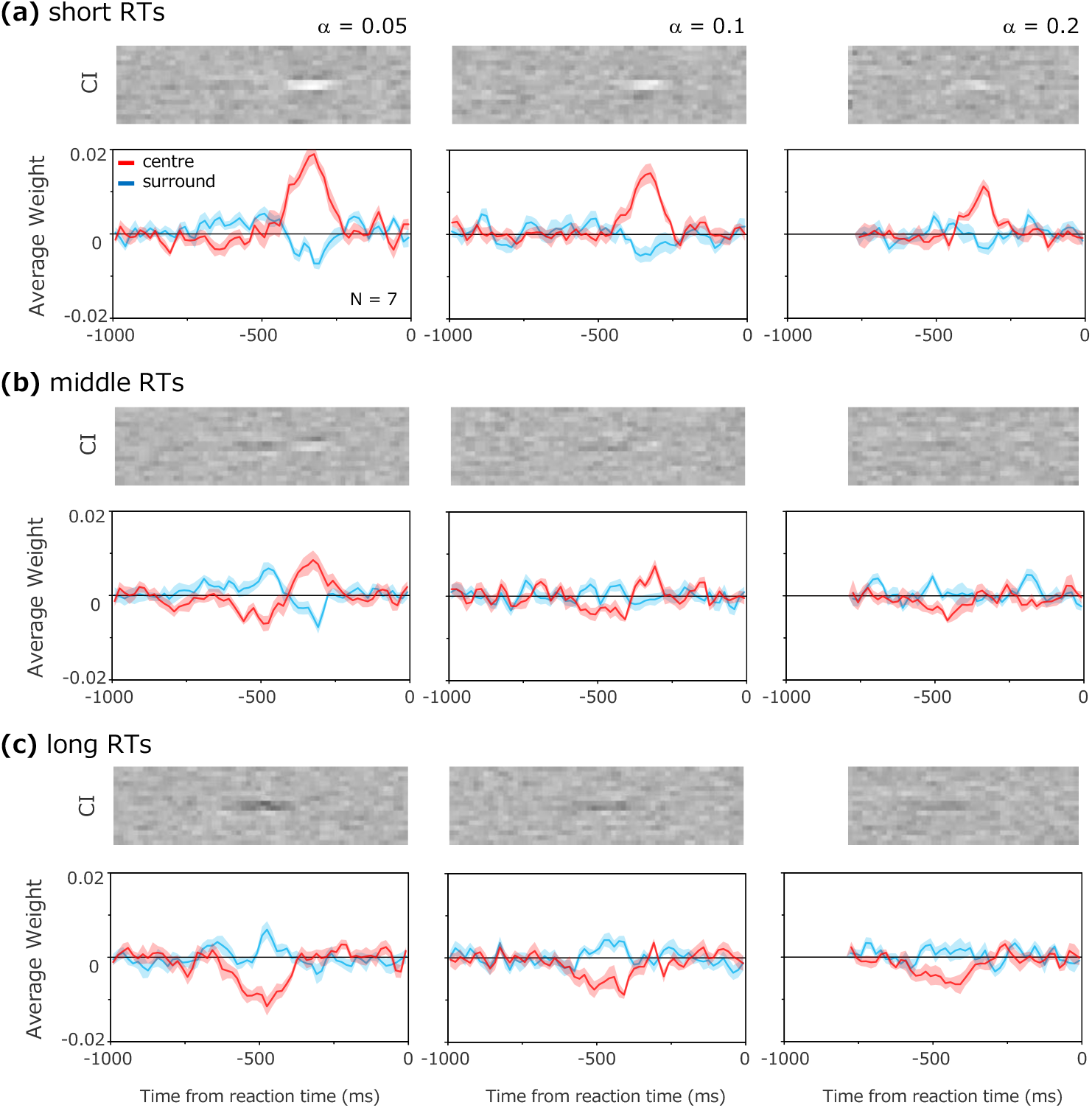
Results for each reaction time group. Results for (a) short reaction times, (b) intermediate reaction times, and (c) long reaction times. Upper panels show the CIs and lower panels show the impact curves.

For each condition, we calculated the average weights during the time epoch from −250 to −350 ms and during the time epoch from −450 to −550 ms. For each time epoch, we performed a three-way-measure ANOVA on the average weights for the position (center and periphery), the contrast increase rate (a = 0.05, 0.1, 0.2), and reaction time group. For the time epoch of −250 to −350 ms, the results show that the main effects of the position (F(1,6) = 269.03, p < 0.001) and reaction time group (F(2,12) = 28.44, p < 0.001) were both significant, and the main effect of the contrast increase rate (F(2,12) = 1.85, p < 0.20) was not significant. The results also show a significant interaction between the position and reaction time group (F(2,12) = 54.025, p < 0.001), between the contrast increase rate and reaction time group (F(4,24) = 4.85, p < 0.01), between the position and contrast increase rate (F(2,12) = 18.18, p < 0.001), and between the position and contrast increase rate and reaction time group (F(4,24) = 5.08, p < 0.005). For the time epoch of −450 to −550 ms, the results show that the main effects of the position (F(1,6) = 114.73, p < 0.001) and reaction time group (F(2,12) = 12.12, p < 0.005) were significant, and the main effect of the contrast increase rate (F(2,12) = 0.25, p < 0.784) was not significant. The results also show a significant interaction between the reaction time group and position (F(2,12) = 7.01, p < 0.01), between the reaction time group and contrast increase rate (F(4,24) = 3.22, p < 0.05), and between the position and contrast increase rate (F(2,12) = 5.90, p < 0.05). There was no significant interaction between the position, contrast increase, and reaction time (F(4,24) = 0.750, p = 0.57).

## 4. Discussion

The present study examined the information utilization strategy adopted in dynamic decision making during stimulus observation in a simple contrast detection task. Applying the classification image method, we calculated the weights of the embedded noise at each time point retrospectively from the reaction time for the target. The resulting CIs indicate that observers responded by utilizing the biphasic luminance change and the central antagonistic spatial contrast before the response. In addition, we found that these spatiotemporal profiles of CIs varied significantly depending on the reaction time.

The complex diversity of the results depending on the reaction time appears to be difficult to understand intuitively. This may seem to indicate that the observers were so flexible that they use different strategies for utilizing information depending on whether they could respond quickly to the target or not. However, we should first consider a simple explanation—that the results are a natural consequence of an interplay between the sensory system and the decision process. Therefore, we tested a simple model consisting of the early visual process (linear filtering model) and the perceptual decision process (drift-diffusion model). As a result, we found that this conservative model with a fixed set of parameters successfully duplicated the human data for all conditions and RT ranges.

### 4.1. Computational model

Figure 6 shows an outline of the model, which is inspired by a previous study on spatiotemporal ensemble perception (Yashiro et al., 2020). The model compares the spatially summarized outputs of the perceptual process, which is approximated using linear spatiotemporal filters, between the two regions. The decision process accumulates the differential signal between the two regions as sensory evidence over time and makes a decision when the evidence reaches a given boundary. The basic structure of the perceptual process follows that of a previous CI study (Neri & Hereger, 2002), and the computation of the decision making follows traditional DDM modeling for a two-alternative forced-choice task (Ratcliff & McKoon, 2008; Gold & Shadlen, 2007; Kiani, Hanks & Shadlen, 2008). Fig. 6 shows each step of the process graphically for the case that a target appears on the left. The calculation of each step is described in detail below.

**Figure 6.**
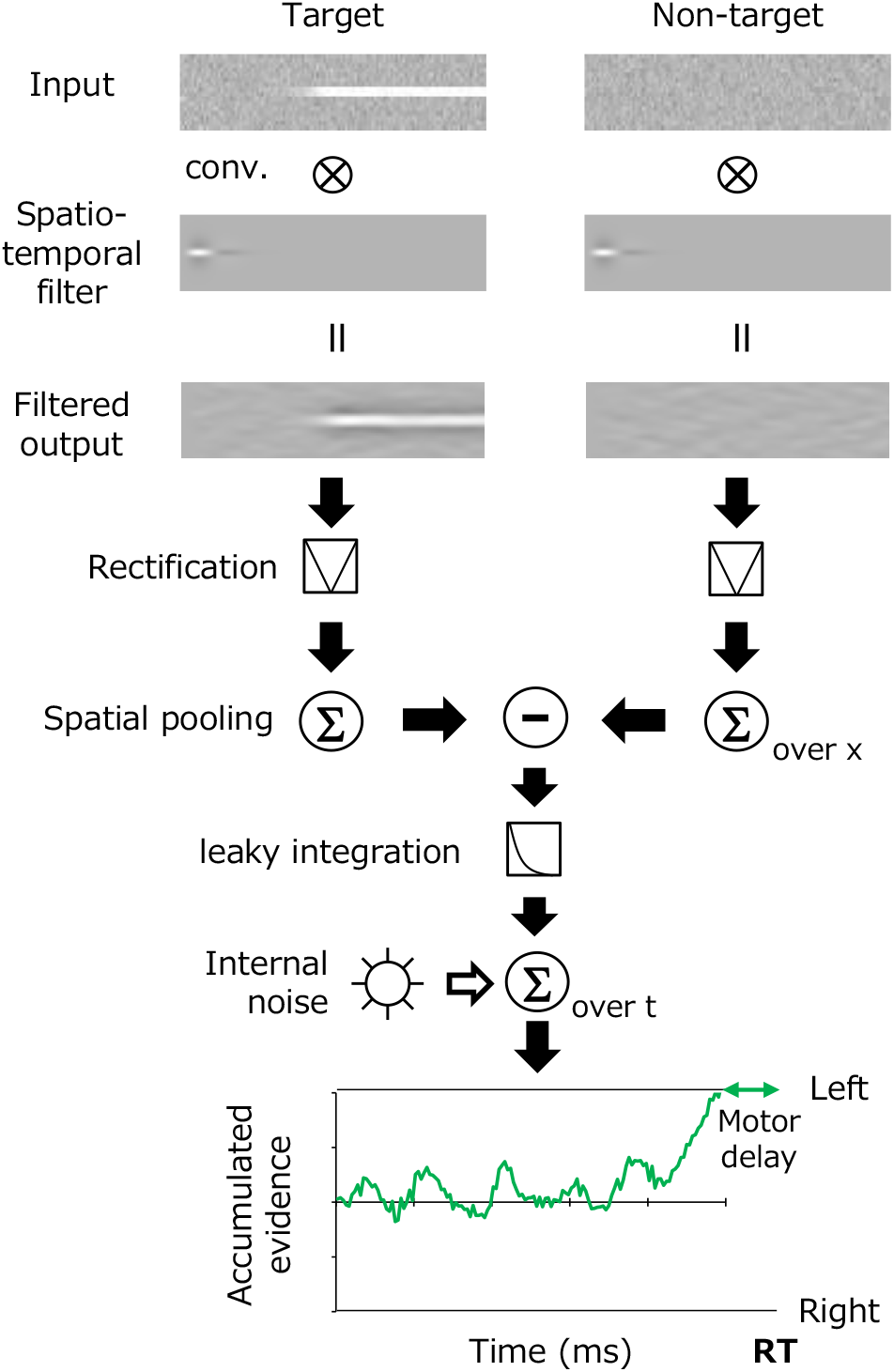
Schematic diagram of a model based on spatiotemporal filtering and the accumulation of sensory evidence.

Following previous studies (Neri & Hereger, 2002), the perceptual system is approximated as a linear spatiotemporal filter, *F_st_(x,t)*, as follows.

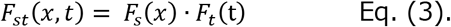

Here, *F_s_(x)* is the spatial filter and *F_t_(t)* is the temporal filter. The spatial filter *F_s_(x)* is given as a DoG function, which has been widely used as a first-order approximation for contrast detectors in the visual system (Enroth-Cugell & Robson, 1966).

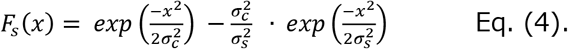

Here, *σ*_c_ is the standard deviation for the central region and *σ*_s_ is the standard deviation for the surrounding region. The temporal filter *F_t_(t)* is given as the following biphasic function (Watson, 1986; Adelson & Bergen, 1985).

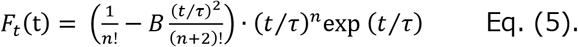

Here, *n* is the number of stages in the time integrator, *τ* is the transient factor, and *B* is a parameter that defines the amplitude ratio of the positive and negative phases.

The response of the perceptual system was obtained by convolving the above spatiotemporal filter *F_st_(x,t)* with the stimulus input *I(x,t)*.

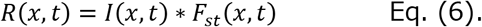

Decisions concerning whether the target presented in the left or right region were made by comparing the spatial sum of the absolute values of the responses in each region between the left and right. Thus, the model observer continually monitored the difference *Δ*R(t) between the left and right responses at time *t* from the stimulus onset. Here, *Δ*R(t) is regarded as the sensory evidence at time *t* in the decision-making model.

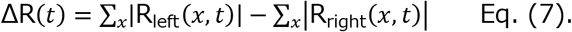

Decisions for targets are based on evidence accumulated over time. However, a number of decision-making studies suggest that sensory evidence decays with time; that is, the evidence weakens as it ages (Usher & McClelland, 2001; Hanks & Summerfield, 2017; Yashiro et al., 2019). This property is practically described as a leaky temporal integration, and it is potentially a product of the adaptive gain control of evidence signals (Cheadle, Wyart, Tsetsos, Myers, Gardelle, Castañón & Summerfield, 2014; Li, Michael, Balaguer, Castañón, Summerfield, 2018). According to these findings, the present modeling assumes that the cumulative evidence *S(T)* at time *T* is given by the following equation, which approximates the noisy leaky integration of *Δ*R(t).

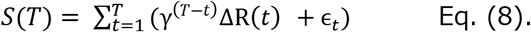

Here, *γ* is the time constant of evidence integration and *ε*_t_ is the internal noise following a normal distribution. The model observer makes a decision about whether the target is on the left or right when *S(T)* exceeds a certain decision boundary, that is, *β*or *−*β** respectively. The observer was assumed to execute a manual response after a constant motor delay of 250 ms from *T*.

In this modeling, the perceptual process part has five parameters: the standard deviations of the spatial filter (*σ*_c_ and *σ*_s_), the number of biphasic temporal filter integration stages (*n*), the time constant (*τ*) and the ratio of positive and negative phases (*B*). The decision-making process part has three parameters: the decision boundary (*β*), the internal noise (*ε*_t_) and the time constant for evidence reduction (*γ*).

### 4.2. Model Simulation

We analyzed the CI and impact curves of the model observer using the image input data that were presented to each observer in the experiment. In the simulations, for all data in the condition of α = 0.05, the model parameters were optimized for each observer to minimize the squared error between the impact curve obtained for the model observer and that of the human observer. To achieve the steady fitting, only the number of integration steps of the biphasic temporal filter (*n*) was fixed to 5, for all model observers.

Figure 7 shows the simulation results. The thick impact curve represents the average of results obtained for the optimized model for each observer, and the light-colored bands represent the ±1 se range of the average for the human observer data. Estimated parameters and the s.e. across model observers were [*σ*_c_, *σ*_s_, *B*, *τ*, *γ*, *β*, *ε*_t_] = [4.15, 17.4, 0.47, 0.793, 0.046, 496.5, 49.6] (s.e. = 0.18, 1.67, 0.07, 0.084, 0.002, 3.58, 2.64). For all values of the contrast increase (α), we find that the model successfully duplicated both the CI and the impact curve of the observers. For the three different reaction time groups (Fig. 7b–d), the model duplicated the observers’ data, reflecting the characteristic differences of RT-dependent CI and impact curves. The root-mean-square error (i.e., difference) between the fitted models and the model observers’ data averaged over all observers was 0.005 (s.e. = 0.0001).

**Figure 7.**
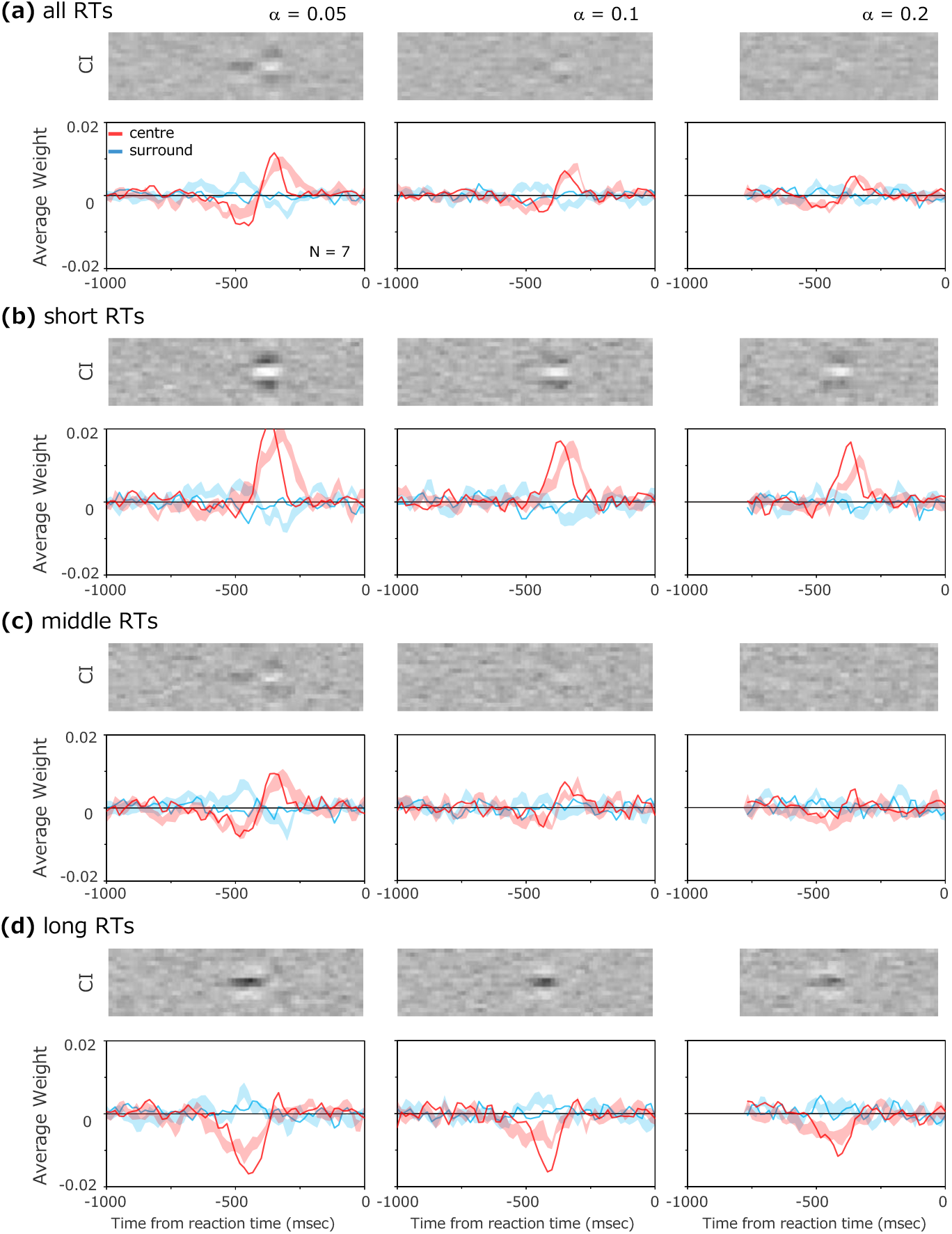
Results of the model simulations. (a) Impact curves and CIs derived from the data for all RTs. (b)–(d) Impact curves and CIs when divided into reaction time groups, with solid lines representing results for model observers and light-colored bands representing the ±1 s.e. range for human data.

To investigate the importance of the functional processes assumed in the model in Fig. 6, we simulated the model without some of the functional processes. We found that (1) the unique shape of the observers’ CIs and impact curves could not be simulated if even one of the parameters of the spatiotemporal filter was omitted and (2) without the leaky integration property being assumed in the decision making, the effect of the early stage of stimulus presentation did not decrease even after a long observation in some RT ranges. On the other hand, modifying the model to accumulate the responses in each domain separately as two pieces of evidence and then calculate those differences, instead of accumulating the differences in responses between the two domains as evidence, did not change the behavior of the model, because the model essentially accumulates evidence linearly (c.f., Bogacz, Brown, Moehlis, Holmes & Cohen, 2006).

The present results support the idea that on-the-view behavioral responses to visual stimuli can be explained by a simple combination of the conventional perceptual model and the standard perceptual decision-making model. This finding may allow us to perform a response-locked reverse correlation analysis of human responses to sensory stimuli during observation, rather than after observation, to explore the characteristics and strategies of human information use in various cognitive tasks. In further investigations, a similar framework may be used to understand the mechanisms for attentional selection and for high-level visual cognition. The present computational model can be used as a baseline account in these investigations.

It should be noted that psychophysical analysis cannot reliably separate the properties of decision making from the low-level perceptual process (Okazawa et al., 2018). Although one can partially overcome this limitation by making full use of various aspects of data, such as by dividing the data into different RT ranges as in the present study, it is difficult to distinguish between some properties such as the latency of the perceptual sensors and the motor delay in the decision process.

## Acknowledgements

This study was supported by the Commissioned Research of NICT(1940101) and JSPS KAKENHI JP20H01782.

## Notes

### Competing Interest Statement

The authors have declared no competing interest.

